# Genome-wide identification of silencers in human cells

**DOI:** 10.1101/2023.06.20.545673

**Authors:** Xiusheng Zhu, Chao Wang, Dashuai Kong, Jing Luo, Biao Deng, Yi Gu, Siyuan Kong, Lei Huang, Yuwen Liu, Yubo Zhang

## Abstract

Transcriptional regulation is a complex process that is controlled by a variety of factors, including enhancers and silencers. Silencers, also known as repressor elements, play a crucial role in the fine-tuning of gene expression by inhibiting or suppressing transcription in the human genome. Although significant progresses have been made, genome-wide silencer research is still in its early stages. Here, we used a genome-wide method called massively parallel reporter assays (MPRAs) to identify silencers in three human cell lines: K562, LNCap, and HEK293T. We identified 739,434, 643,484, and 491,952 silencer regions in these cell lines, respectively. We found that most of the silencers we identified had inhibitory activity and significantly enriched inhibitory motifs. These results confirm that silencers are ubiquitous in the human genome and play an important role in regulating gene expression. Therefore, our study provides a general strategy for genome-wide functional identification of silencer elements. This information could be used to better understand the mechanisms of gene regulation and to develop new therapeutic strategies for diseases that are caused by dysregulation of gene expression.

## Introduction

Cis-regulatory elements within the non-coding regions of the genome play a pivotal role in governing gene transcription and influencing diverse biological traits. In recent years, Roadmap Epigenomics and Encyclopedia of DNA Elements (ENCODE) have made significant strides in the annotation of regulatory elements and shedding light on non-coding regions[1-4]. Substantial progress has been made in unraveling the intricacies of transcriptional regulatory elements, such as enhancers and promoters. Understanding the mechanisms underlying these regulatory elements not only enhances our knowledge of disease etiology but also holds promise for precise gene therapies[5].

Nevertheless, the current focus of research on regulatory elements has primarily centered around enhancers, while silencers, another crucial class of regulatory elements, have received relatively limited attention.In 1985, Brand et al.identified silencer in yeast for the first time,, marking a seminal moment in the field[6]. Unfortunately, subsequent investigations into silencers experienced a decline in interest, with only a handful of specific silencers, such as those associated with the human thymocyte CD4 gene[7,8] and the neuron restrictive silencer element (NRSE)[9], being studied in depth.Recent reports have highlighted the development of high-throughput screening methods aimed at identifying silencers. For instance, Pang and Snyder introduced the “ReSE” method, which successfully identified over 2,000 silencers, representing approximately 1% of the human genome[10]. Jayavelu et al.[11] employed subtraction analysis and MPRA to identify silent sub-elements within the genome, subsequently employing algorithms to predict silent sub-elements in both human and mouse genes. However, none of these studies have conducted comprehensive functional validation of silencers at the genome-wide level. Prediction of silencers based on specific histone modifications or binding proteins, representing an alternative approach, has been explored by Nagn et al., who described silencers and their chromatin interactions in mouse embryonic stem cells using PRC2 ChIA-PET data[12] Cai et al. identified silencers within the human genome using ChIP-seq data of H3K27me3[13]. However, the characteristics of silencers do not consistently align with these histone modifications, leading to inherent biases in silencer screening..

While STARR-seq has proven effective in identifying enhancers through transcription level changes, experimental methods for genome-wide silencer analysis remain elusive[14]. The construction of screening libraries, particularly for the vast human genome, poses significant challenges. Additionally, existing enhancer-focused algorithms for STARR-seq and conventional silencer analysis methods are ill-suited for comprehensive genome-wide silencer investigations[15-19]. A recently published method called CRADLE (correcting reads and analysis of differentially active elements) has emerged, offering the potential to identify silencers within STARR-seq data, ensuring accurate genome-wide silencer data identification[20].

In this study, we present the construction of an unprecedented whole-genome screening library dedicated to silencers. By leveraging human cell lines K562, LNCap, and HEK 293T as research models, we have experimentally obtained a comprehensive dataset of human whole-genome silencers, marking a significant breakthrough in the field.

## Results

### Identification of human genome-wide silencer

To facilitate the screening of silencers using the STARR-seq vector, we optimized the vector design. Specifically, we replaced the original SCP1 weak promoter with the hPGK strong promoter, which has been previously utilized for silencer screening[10]. Additionally, we introduced the GFP gene downstream of the promoter to enable visualization of cell transfection efficiency (Figure 1A). Subsequently, we identified silencers within K562, LNCap, and HEK 293T cells. Briefly (see the “Methods” section for detailed procedures), genomic fragments were sonicated to obtain ∼200 bp fragments, which were then ligated to the screening vector to construct an input library. High-throughput sequencing of the input library DNA revealed that it covered approximately 91.4% of the human genome region, with an average coverage of 30.1x coverage of 30.1x (Figure 1B). This unprecedented coverage and depth provided a prerequisite for the genome-wide identification of silencers. Following the STARR-seq output library construction method, the input library plasmid was transfected into cells. After 24 hours, cellular RNA was collected and reverse transcribed into cDNA. Subsequently, the cDNA served as a template for PCR amplification prior to Illumina high-throughput sequencing. Each cell type was subjected to three independent biological replicates, and the results demonstrated high correlation coefficients among the replicates (Figure 2 A, B, C), confirming the reproducibility of the technology. Finally, using the statistical method for identifying silent subregions (see the “Methods” section for details), we identified a total of 739,434,643,484,491,952 silent subregions in K562, LNCap, and HEK 293T cells, respectively. These silent subregions exhibited robust repeatability (Figure 3 A, B, C),rendering them suitable for downstream signal analysis.

**Figure 1:**
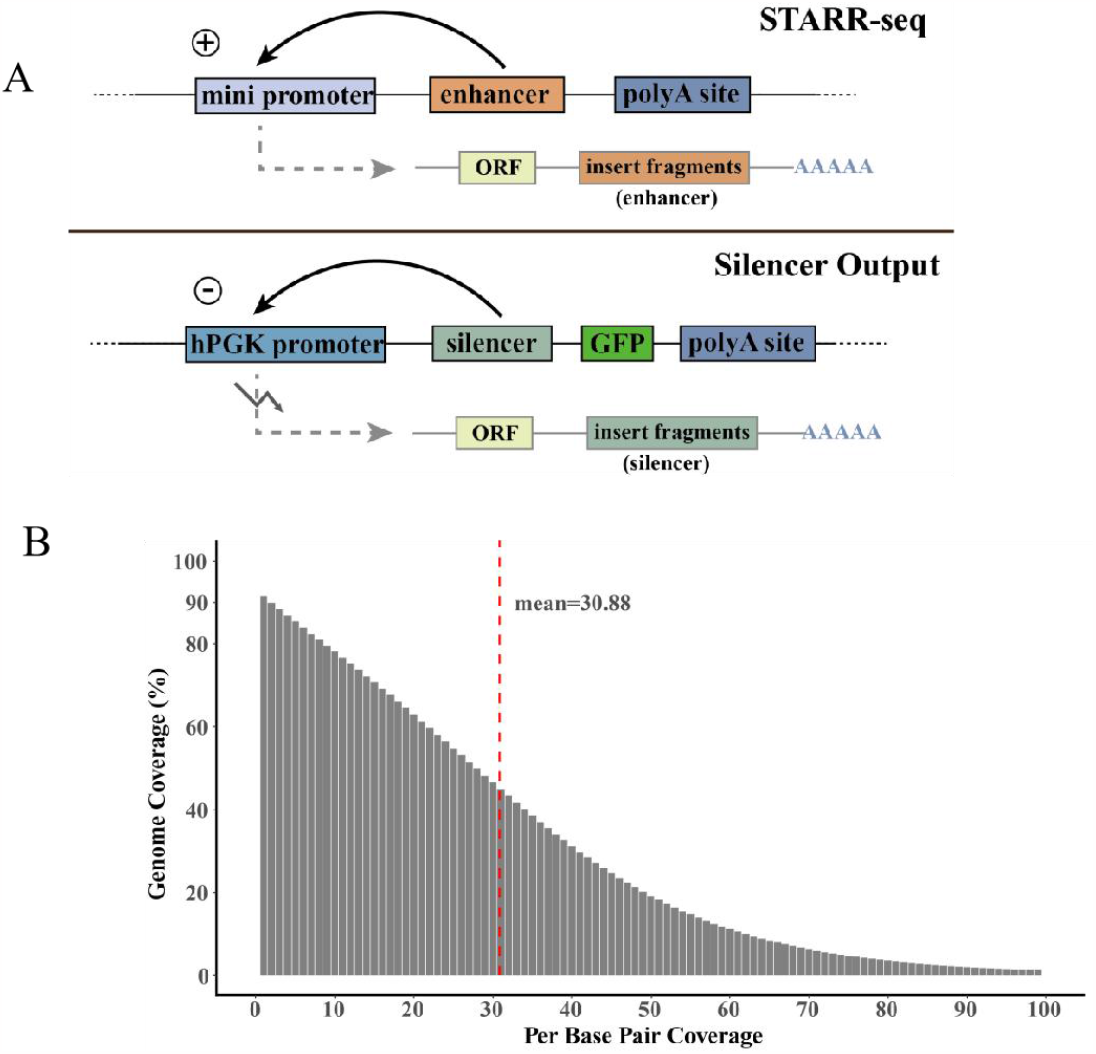
The complexity of silencer screening system and input library. A: Compared with STARR-seq, we replaced the promoter with a strong promoter hPGK and inserted the GFP reporter gene to detect the transfection efficiency. B: The genomic fragments were disrupted into about 300 bp fragments and inserted into the plasmid vector (input library). The fragments inserted into the plasmid vector covered 91.4 % of the human genome, with an average depth of 30.88.

**Figure 2:**
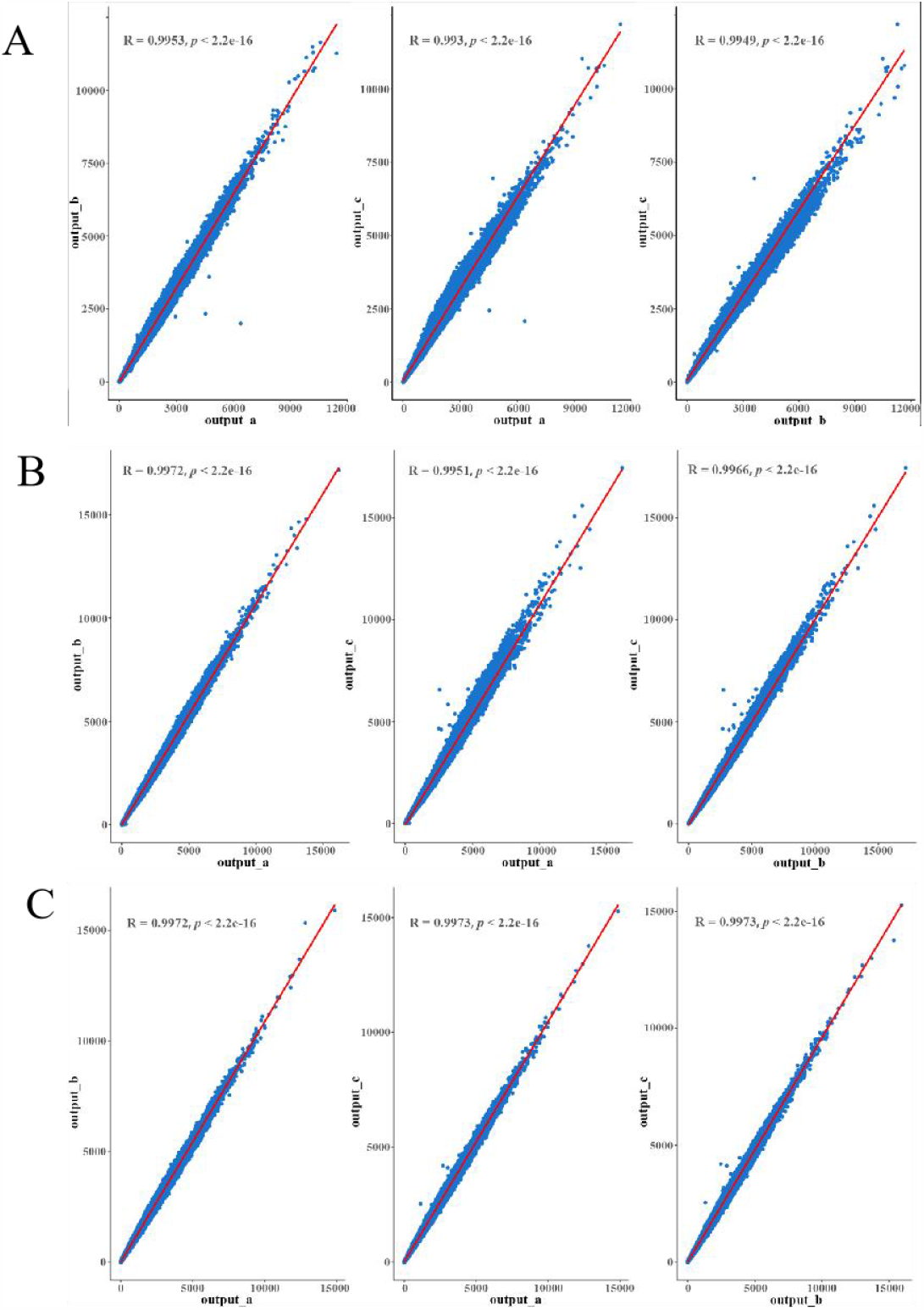
Correlation analysis of output library in HEK 293T, LNCap, K562 cells. A: The correlation analysis of three output libraries in HEK 293T cells showed that the R value was above 0.99. B: The correlation analysis of three output libraries in LNCap cells showed that the R value was above 0.99. C: The correlation analysis of three output libraries in K562 cells showed that the R value was above 0.99.

**Figure 3:**
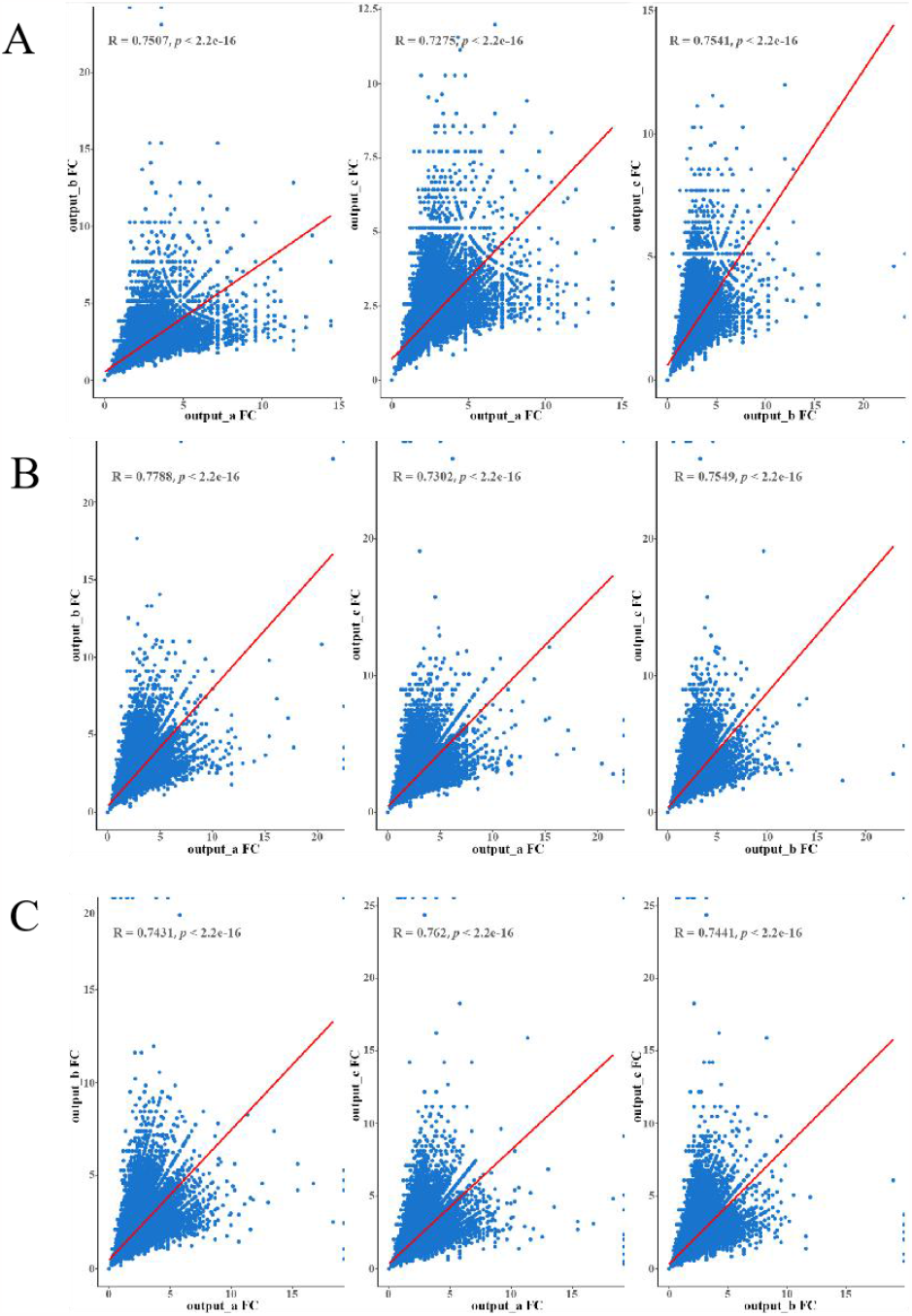
The correlation between the silencer activity of HEK 293T, LNCap and K562 cells in different output libraries was evaluated. A: The correlation of silencer activity in HEK 293T cells between different output libraries was evaluated, and the R value was above 0.7. B: The correlation of silencer activity in LNCap cells between different output libraries was evaluated, and the R value was above 0.7. C: The correlation of silencer activity in K562 cells between different output libraries was evaluated, and the R value was above 0.7.

Next, we examined the activity distribution of these identified silencers. The analysis revealed that the activity of these silencers predominantly fell within the range of 1-2 (Figure 4A), indicating lower activity levels compared to enhancers[21]. This discrepancy may arise from differences in the screening systems employed or the inherent mechanisms of silencers themselves. We further explored the genomic distribution of these silencers and observed that they were predominantly located in distal intergenic regions, intronic regions, and promoter regions in the distal intergenic regions, intron regions and promoter regions (Figure 4B). To assess whether these silent subregions exhibited enrichment of inhibitory transcription factor motifs, we employed SeqPos21 to perform motif analysis[22]. The results demonstrated that the repressor motif REST was prominently enriched among the top motifs identified in each cell (Figure 5A, B, C). In addition, some motifs such as CTCF and SP family motifs were found to be enriched in both cells (Figure 5A, B, C), suggesting t their potential importance in transcriptional inhibition. Finally, to validate the functionality of the identified silent subregions, we randomly selected 30 of them for dual luciferase reporter assay. Remarkably, 29 of the selected subregions exhibited varying degrees of inhibitory activity, thus confirming the accuracy of our identified silent subregions.

**Figure 4:**
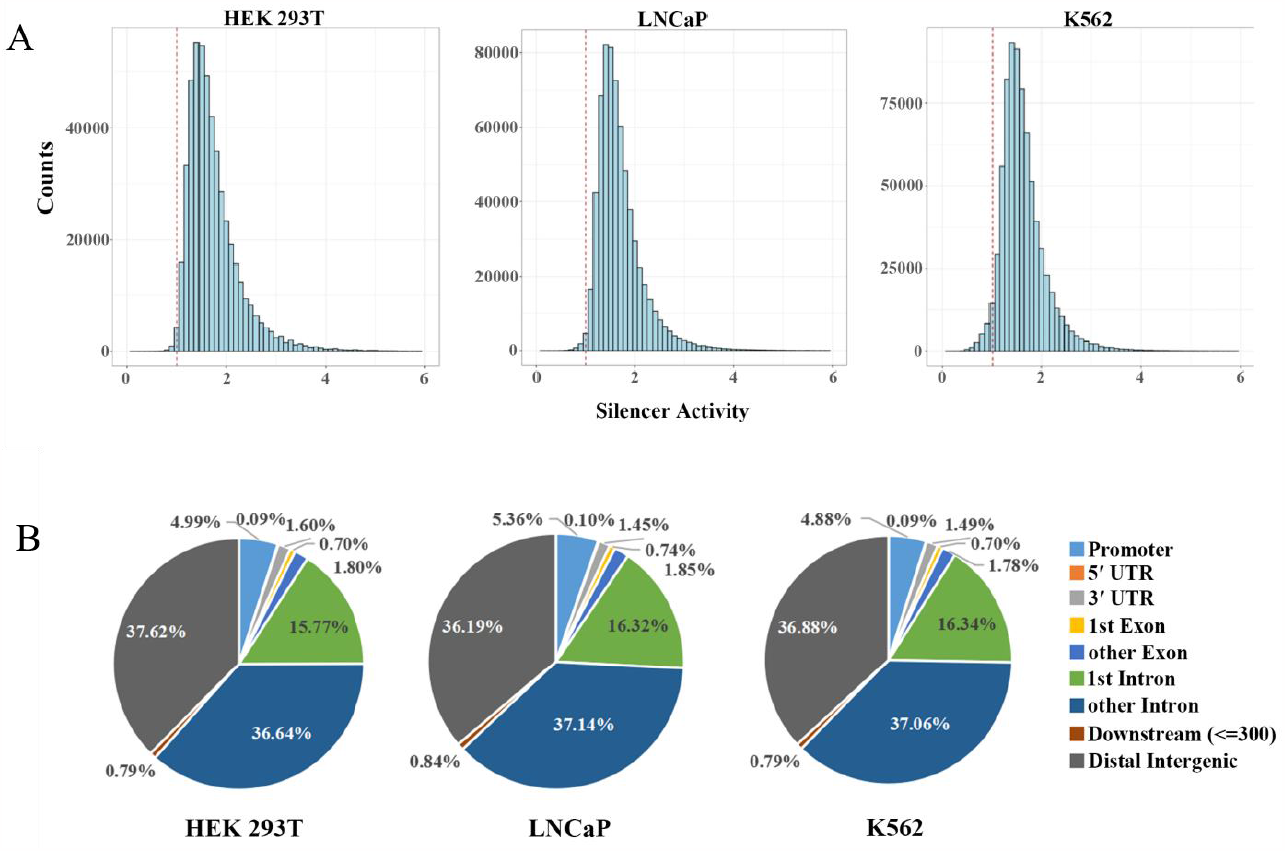
HEK 293T, LNCap and K562 cells were used to identify the activity distribution and genomic region distribution of silencers. A: The distribution of silencer activity in HEK 293T, LNCap and K562 cells is mostly concentrated between 1-2. B: Genome-wide distribution of silencers in HEK 293T, LNCap and K562 cells.

**Figure 5:**
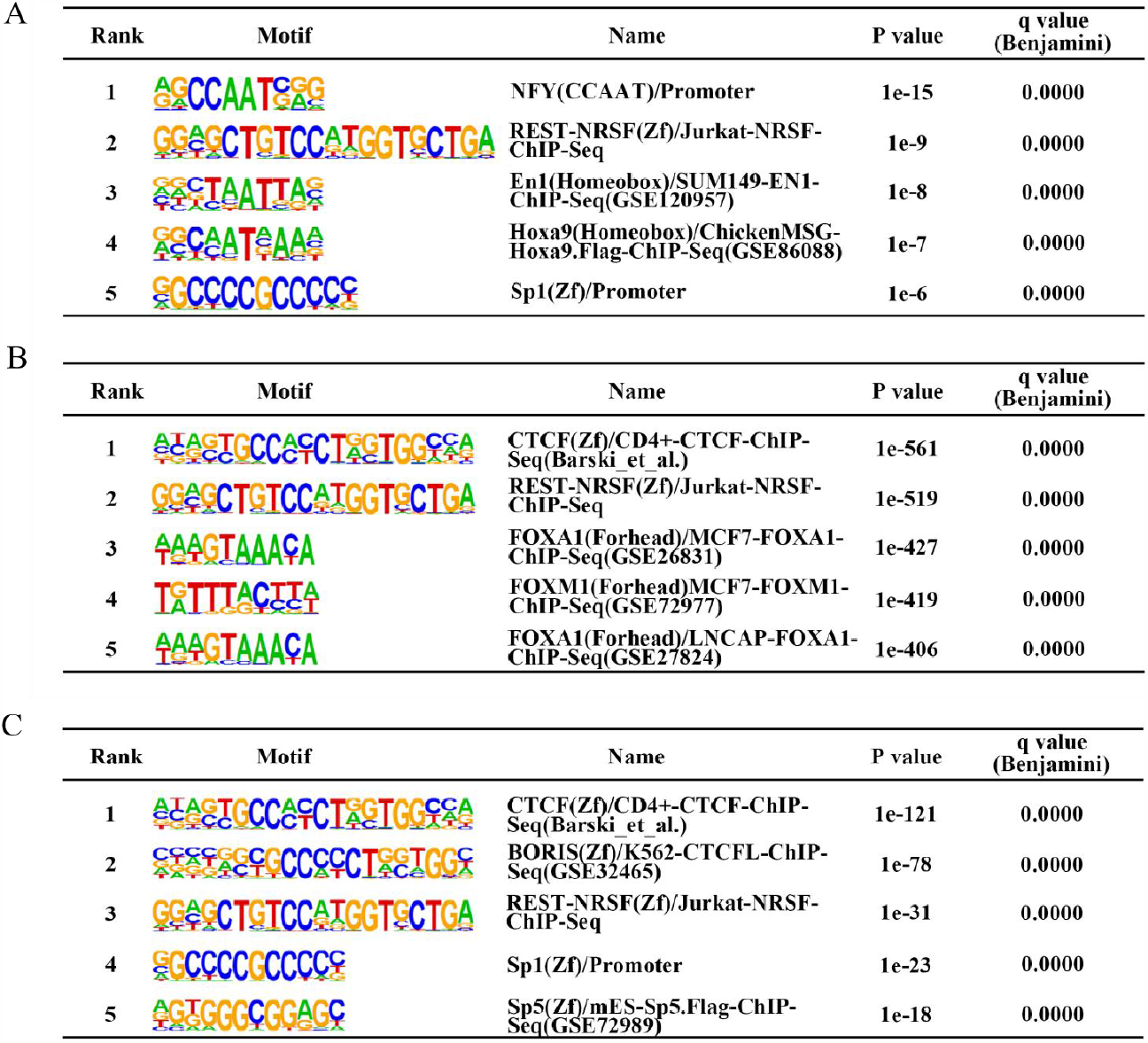
Motif enrichment analysis of silencers identified by HEK 293T, LNCap and K562 cells. A: Motif enrichment analysis of silencer regions in HEK 293T. B: Motif enrichment analysis of silencer regions in LNCap. C: Motif enrichment analysis of silencer regions in K562.

**Figure 6:**
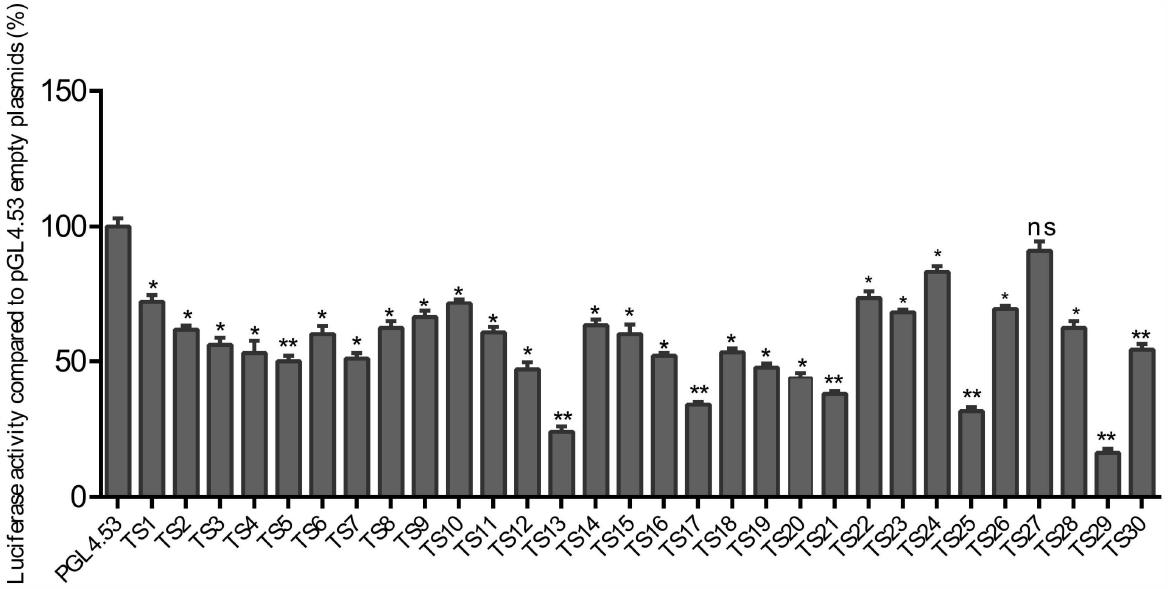
Luciferase assays to determine the repressive activity of silencers identified from HEK 293T, LNCap and K562 cells. The empty PGL 4.53 plasmid was the control group. The y axis represents the percentage of luciferase activity compared to pGL4.53 empty plasmids in the respective cells (3 biological replicates per sample; bars show mean value±s.e.m. *: p<0.05, **:p<0.01, ns: no significant difference, calculated using t-test).

## Discussion

A significant portion, approximately 98%, of the human genome lacks coding function but contains cis-regulatory elements that play a crucial role in gene expression regulation. However, current research primarily focuses on enhancers and promoters, possibly due to their positive impact on gene expression regulation. In contrast, the study of silencers, which exert inhibitory effects on gene expression, has been limited, primarily due to the lack of genome-wide data and understanding of their mechanisms of action.

In recent years, several studies have reported the identification of silencers through high-throughput methods. However, most of these studies have not achieved genome-wide coverage. Computational simulation methods can provide genome-wide data on silent subregions; however, they heavily rely on the quality of the training data and often lack experimental validation. Furthermore, unlike enhancers that exhibit specific histone modifications, such as H3K4me1 and H3K27ac[21], enabling their identification through ChIP-seq, silencers lack specific histone modifications. Consequently, using specific chromatin features to predict silencers may lead to numerous false negatives.

To address these challenges and from a functional perspective, we experimentally identified approximately 1.9 million silencers in human cells. Subsequent experiments confirmed that the majority of these identified silent subsequences exhibited inhibitory activity. Previous reports predominantly focused on silencers identified in open chromatin regions; however, the majority of the genome exists in closed chromatin regions. Therefore, it remains unclear whether the inhibitory activity of silencers in closed chromatin regions is influenced by the chromatin environment. Additionally, it is essential to investigate whether silencers, similar to enhancers, form interaction loops with promoters to regulate gene expression. Furthermore, exploring whether silencers possess specific histone modifications and assessing their impact on biological phenotypes necessitate comprehensive genome-wide silencer data for further analysis.

In summary, our study successfully constructed a genome-wide map of silencers in three distinct human cell types, providing high-quality data for analyzing the mechanisms underlying silencer function. This precise genome annotation of silencers contributes to a deeper understanding of gene regulation networks, shedding light on the intricate interplay between regulatory elements and gene expression.

## Methods

### Input library construction

The random input DNA library is mainly constructed in three steps (insert fragment construction, linearized vector construction and seamless assembly clone). The genome was randomly disrupted into about 300 bp DNA fragments by using an ultrasonic disruptor, and the library construction kit (Roche; KK8514) linked the homologous arm and the double-ended Illumina adapter to both ends of the fragmented DNA to construct an insertion fragment. AgeI-HF enzyme (NEB; r3552L) and SalI-HF enzyme (NEB; r3138 S) digested hPGK-STARR-seq vector, and then used Gibson homologous recombination enzyme (NEB; e2611L). The inserted fragment was connected with the linearized vector by homologous recombination. Sanger sequencing was performed to check the quality of library clones. Sixty homologous recombination reaction systems were prepared and DNA purification kit (ZYMO ; d4013), and the purified product was transformed into Thermo Fisher Scientific (Thermo Fisher Scientific ; c6400). The transformed competent cells were transferred to 15 L LB medium for amplification. When the OD value of the bacterial solution was 1.0-1.2, according to the kit (QIAGEN ; cat 12362). PCR amplification was performed with 100 ng plasmid as template 98°C 45 s ; a total of 20 PCR reactions were prepared (98°C 15 s, 60°C 30 s, 72°C 30 s amplification 20 cycles). The product was recovered by 2 % agarose gel electrophoresis for 40 min (120V, 130mA) and subjected to high-throughput sequencing.

### Cell culture and transfection

HEK 293T cells were inoculated with 10 % FBS (Gibco; cat 10437028) and 1% penicillin streptomycin P/S (Sigma; cat P0781) in DMEM (Thermo Fisher; cat 10566024), K562 cells were inoculated with 10% FBS (Gibco; cat 10437028) and 1 % penicillin streptomycin P/S (Sigma ; LNCap cells were seeded in IMDM (iCell, Cat0008) medium containing 10 % FBS (Gibco; cat 10437028) and 1 % penicillin streptomycin P/S (Sigma; cat P0781) RPMI-1640 (Gibc ; cat11875) was cultured in a cell incubator at 37 °C. When the cell growth confluence was about 80%, Nucleofection electroporator (Lonza; aAF-1002X) was transfected into the input library (1ug DNA /10^6^ cells). The cells after 24 hours of transfection were collected to construct the output library.

### Construction of output library

The cells transfected with the input library were collected for RNA extraction using TURBOTM DNase (Thermo Fisher Scientific; aM2238) using Dynabeads Oligo TM (dT)25 (Thermo Fisher Scientific ; 61002). SuperScript TM IV Reverse Transcriptase (Thermo Fisher Scientific; cat.18090200) and 2M specific primers (5’ - CAAACTCATCAATGTATCTTATCATG-3’) were used for reverse transcription of the enriched mRNA to generate a cDNA product, and then AmbionTM RNase H was used to remove the residual RNA in the cDNA product. After purification of the cDNA product, the HFi high fidelity enzyme ReadyMix (Roche ; kK2601) for PCR amplification (98 °C 45s; after20cycles of amplification at 98 °C for 15s, 60°C for 30s, and 72°C for 30 sec, the products were recovered by 2% agarose gel electrophoresis for 40min (120V, 130mA) to construct the output library for high-throughput sequencing.

### Dual-luciferase assay

To synthesize silent subsequences to be verified (GENEWIZ, China). These sequences were inserted into the hPGK promoter of the luciferase plasmid pGL4.53 (Promega) using Gibson assembly. Then the reporter vector and the pGL4.53 vector inserted with the silent subsequence were co-transfected into the cells. According to the manufacturer’s plan, the Dual-Glo luciferase assay kit from Promega is used for luciferase analysis. The original luciferase plasmid without any insertion sequence was used as a control. All luciferase assays were three independent transfections performed on different dates.

### Library Quality Control

The input library and output library after Illumina sequencing were subjected to quality control. The TrimGalore software was used to perform quality control on the offline data, including connector removal, low-quality base and short sequence removal (-q20-length 25). For the clean data generated by quality control, the Bowtie2 software was used to compare the clean data with the human reference genome hg38 to generate a BAM file. Sambamba software was used to remove PCR repeated reads, and then SAMtools software was used to extract the BAM file of double-end reasonable alignment reads.

### Genome coverage analysis of the library

The genomecov command of BEDtools was used to calculate the number of coverages of reads in each base of the genome in the input library. The number of bases with a coverage of more than 1 was counted in the result file, which was divided by the total number of bases in the genome.

### Genome-wide identification of silent sub-elements

Based on the BAM file of the input and output libraries containing only reasonably aligned reads at both ends, it is converted into a bw file (--normalizeUsing None) by the bamCoverage command, and then the CRADLE software is used to identify the silent sub-region of the whole genome. The first step is to use the correctBias _ stored function to perform technical bias correction on read counts (shear, PCR, mappability, G-quadruplex), and then use the callPeak command to identify the DNA region with regulatory activity and inhibitory effect of the whole gene (see https://pypi.org/project/CRADLE/), for detailed screening of DNA regions with inhibitory effects as silent sub-regions.

### Library repeatability evaluation

In order to evaluate the repeatability between different libraries, we adopted two methods to evaluate the repeatability of the library. One is to evaluate the consistency of the distribution of reads in the whole genome, and the other is to evaluate the consistency of the activity of the whole genome silencer. Two evaluation methods are implemented by Deeptools software. The evaluation of genome-wide reads distribution consistency uses the multiBamSummary bins command to divide the genome into 10 kb bins, count the count number under each bin, and then perform library depth correction, and then use plotCorrelation to draw the correlation between libraries. The correlation evaluation of genome-wide silencer activity between different libraries was performed by using the multiBamSummary BED-file command to count the counts of silencer regions in different libraries. After deep correction of library sequencing, the Input RPM / output RPM value was used to represent the activity intensity of each silencer. Based on the activity intensity of all silencers in different libraries, the correlation analysis and visual display of silencer activity between libraries were performed using R ggplot2 and ggpbur packages.

## Author contributions

Y.Z.,X.Z. and Y.L. conduct and design the experiments. X.Z., D.K performed the experiments. S.K. and B.D help X.Z performed the experiments. The work of bioinformatics analysis is completed by C.W. and J.L. The manuscript is written by X.Z. and Y.Z. All authors read and approved the final manuscript.

